# Ultra-sensitive proteome profiling of FACS-isolated cell populations by data-independent acquisition-MS: Application to human hematopoietic stem and progenitor cells

**DOI:** 10.1101/371161

**Authors:** Sabine Amon, Fabienne Meier-Abt, Ludovic C Gillet, Slavica Dimitrieva, Alexandre PA Theocharides, Markus G Manz, Ruedi Aebersold

## Abstract

Physiological processes in multicellular organisms depend on the function and interactions of a multitude of specialized cell types operating in context. Fluorescence-activated cell sorting (FACS) provides a powerful tool to determine the cell type composition of complex mixtures and to purify highly homogeneous cell populations using a small number of differentially expressed marker proteins. These populations can be further characterized, e.g. by phenotypic or molecular analyses.

We describe an ultra-sensitive mass spectrometric method for the robust quantitative and reproducible proteomic analysis of cohorts of FACS-isolated cell samples. It uses a minimum of post-sorting sample processing steps prior to data-independent acquisition MS on a current generation Orbitrap hybrid mass spectrometer. The method provides highly accurate and reproducible quantitative proteome profiles across the cohort with an average coefficient of variance <15% from as little as 150 ng of total peptide mass. We quantified the proteome of 25,000 sorted human hematopoietic stem/multipotent progenitor cell and three committed progenitor cell subpopulations (common myeloid progenitors, megakaryocyte-erythrocyte progenitors and granulocyte-macrophage progenitors) isolated from five healthy donors. On average, 5,851 protein groups were identified per sample. After stringent filtering, a subset of 4,131 protein groups (≥2 peptides) was used for differential comparison across the 20 samples, defining unique proteomic signatures for each cell type tested. A comparison of proteomic and transcriptomic profiles of the four cell types indicated hematopoietic stem/multipotent progenitor cell-specific divergent regulation of biochemical processes that are essential for maintaining stemness and were detected at the proteome rather than the transcriptome level.

The technology supports the generation of extensive and accurate quantitative proteomic profiles from low numbers of FACS-purified cells providing new information about the biochemical state of the analyzed cell types that is essential for basic and translational research.

## INTRODUCTION

In multicellular organisms, normal physiological functions and pathophysiological mechanisms are the result of the interplay of multiple cell types at various stages of differentiation. A prototypic example is the mammalian hematopoietic system where hematopoietic stem/multipotent progenitor cells (HSCs/MPPs) can differentiate into various functionally divergent cell lineages including the downstream formation of common myeloid progenitors (CMPs), megakaryocyte-erythrocyte progenitors (MEPs) or granulocyte-macrophage progenitors (GMPs) ^1,2^. When this process is altered, e.g. upon genetic or epigenetic changes in HSCs, abnormal (pre)leukemic stem cell subpopulations may form, eventually resulting in clonal hematopoiesis and in the onset of acute myeloid leukemia ^3-5^. To gain insight into the biochemical changes underlying cellular differentiation and to unravel factors involved in the early development of malignant hematopoietic diseases, highly refined analysis of the different subpopulations of the hematopoietic cell system is critically needed ^6^.

Fluorescence-activating cell sorting (FACS) is an established technology for the characterization of cell types. FACS operates by monitoring the differential quantity of few specific antigens at the surface of single cells, thus determining the cell type composition of a sample and supporting the isolation of live, homogeneous populations for further phenotypic or molecular analyses. The hematopoietic cell types mentioned above are characterized by specific expression patterns of the cluster of differentiation molecules CD34, CD38, CD123, CD45RA and CD10 ^7-9^. For many clinically relevant samples, no more than few thousand cells per subpopulation can be obtained with reasonable effort after FACS from a person. For example, the preparation of 25,000 sorted human HSCs requires up to four liters of steady-state blood or a leukapheresis procedure following hematopoietic stem and progenitor cell (HSPC) mobilization, making further upscaling difficult.

Whereas a few thousand cells are routinely analyzed by modern imaging and genomic profiling technologies ^1,2,8-11^, proteomic measurement, particularly the reproducible measurement across sample cohorts, has remained challenging due to larger sample requirement of prevailing mass spectrometric methods. Proteomic analysis of FACS-isolated cells has nonetheless been reported in several studies. Most of those focused on optimizing specific technical parts of the workflow, such as the cell sorting step itself ^12^, sample preparation ^13,14^ or sample fractionation ^15^. Others used 400,000 cells as starting material which restricted the scope of the analyses to large pools of murine samples ^16^ or *in vitro* model systems. To our knowledge, no systematic assessment of the reproducibility or consistency of the proteomic results of sorted cells has been performed, other than comparing protein identification numbers. It is therefore evident that the robustness of the proteomic analysis of such samples should be of high concern because it is not always possible to obtain replicates for many clinically relevant samples.

Here, we present an integrated workflow for the high-coverage, quantitative proteome profiling of sorted cells that uses data independent acquisition (DIA)-MS on the Orbitrap Lumos platform. DIA-MS is a massively-parallel-in-time acquisition method of fragment ion mass spectra of all detectable precursors in a sample. It provides a complete, yet convoluted, quantitative fragment ion map record of a sample ^17^. Peptide-centric analysis ^18,19^ of DIA datasets results in quantitative peptide matrices ^17^ of sufficient consistency and reproducibility to support cross sample label-free proteomic comparisons of large sample cohorts. To date, DIA studies on hybrid quadrupole-time-of-flight (QqTOF) ^18,20,21^ or Orbitrap ^22,23^ platforms used microgram amounts of total peptide mass for analysis (and even larger amounts of actually processed starting material), a quantity that is one to two orders of magnitude above the quantity achievable by FACS isolation of rare cell types.

With an integrated sample preparation strategy and thorough optimization of the MS acquisition scheme, we established a method that reproducibly identifies and quantifies nearly 6,000 protein groups with a median coefficient of variance (CV) of 9% for 125 ng of HEK293 tryptic peptides. This unprecedented performance was used to profile 25,000 cells from sorted highly enriched HSC/MPP, CMP, MEP and GMP subpopulations from five genetically different donors. The method consistently quantified 4,131 protein groups across all samples and identified factors and biochemical pathways involved in quiescence, stemness maintenance and cell differentiation.

## RESULTS

### Optimization of DIA-MS for small sample loads

Most DIA-MS applications reported so far used 0.5-2 μg of total peptide mass per injection ^18, 20-23^. Therefore, we first optimized the acquisition scheme on an Orbitrap Fusion Lumos mass spectrometer to extend the application of DIA-MS to lower sample quantities while minimizing attrition of quantified proteins.

The instrument platforms used for DIA-MS differ fundamentally in the way they transmit ions and acquire mass spectra: QqTOF instruments are typically operated with fixed tandem mass spectra (MS2) acquisition times and therefore duty cycles, because ions are transmitted to the detector without any accumulation step, offering relatively few parameters for fine tuning. In contrast, newer generation instruments equipped with Orbitrap analyzers allow the optimization of data acquisition at several stages, e.g. by varying the scan time (resolution is directly proportional to the detection time of the transient), by optimizing the number of accumulated target ions and/or the maximum accumulation (injection) time prior to the detection event, and finally by executing accumulation and detection steps in parallel, as implemented on the Lumos instrument. Depending on the intended application and the available sample amounts, optimization of these parameters may yield substantial gains in performance.

We therefore studied the dependency of fill times to reach a desired ion population on the amount of tryptic peptides from a HEK293 cell lysate loaded on column. We used a 40 DIA isolation window scheme spanning 400-1000 *m/z* that showed optimal identification results for 500 ng peptide load in combination with a 2 h long chromatographic gradient (data not shown). The data confirm (Supplementary Fig. 1) that, as expected, the time required to reach the target value of accumulated ions varied considerably per scan. For the peptide-rich regions of the retention time (RT) vs. *m/z* graph, specifically between 400-800 *m/z* and a RT range of 10-80 min, the target was reached within a few ms on average at the highest sample loads. Thus, the accumulation time was much lower than the actual scan time of 64 ms (to reach 30,000 resolution at *m/z* 200). To optimize the use of ions within the time constraint of each Orbitrap scan event (ions for the next scan are accumulated in parallel to the fragment ion acquisition of the current scan), we selected a scheme that took advantage of a median injection time of 30-50 ms (Fig. 1a). At 250 ng sample load this already corresponded to a six- to nine-fold increase of the fill time compared to a 2 μg sample load (Supplementary Fig. 1b). To assess the performance limits of the optimized acquisition scheme, we compared the number of protein identifications of a standard data-dependent acquisition (DDA) method to the optimized DIA protocol with triplicate injections of a dilution series (in steps of ½) of HEK293 peptides ranging from 2 μg to 3.9 ng. The results indicate that the DIA mode systematically identified a higher number of peptide precursors and proteins than DDA at all sample loads (Fig. 1b and Supplementary Fig. 2). The average number of identified protein groups decreased by only 12% (from 7,406 to 6,472 applying a precursor Q- value cutoff of 0.01 (see Online Methods)) in DIA when reducing the sample amount from 2 μg to 125 ng of peptide mass on column. For the same concentration range, the fraction of protein groups consistently identified in all three injections remained above 85% in DIA (Supplementary Fig. 3a-b, Supplementary Table 1) and >98% of protein groups identified at low sample loads were also found with the highest loads (Supplementary Fig. 3c).

**Figure 1.**
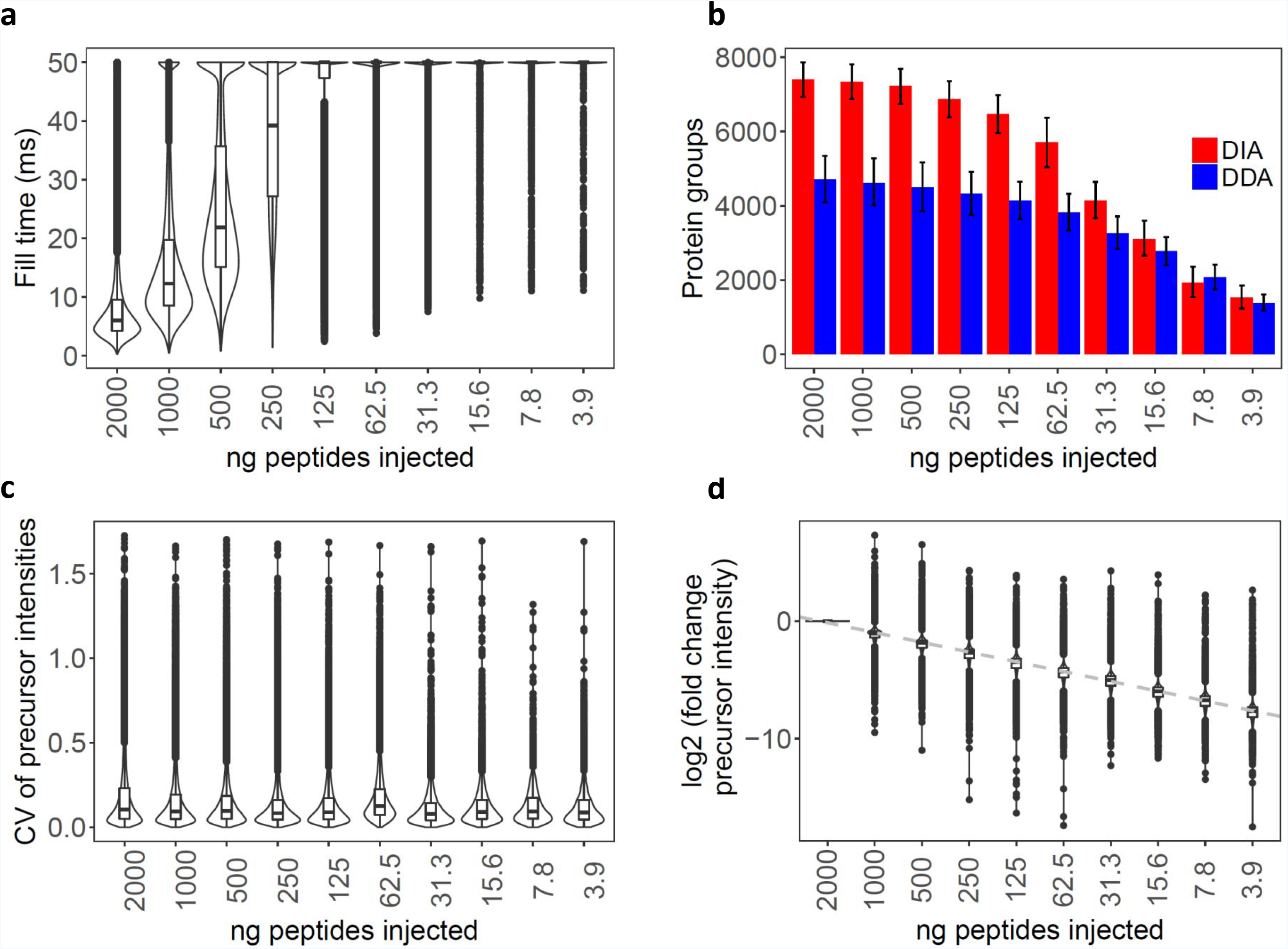
**a.** Distribution of the fill times for the peptide-rich regions of retention time (RT) vs. *m/z* diagram between 400-800 m/z and RT 10-80 min for decreasing loads of HEK293 tryptic peptides (see also Supplementary Fig 1.a). **b.** Number of proteins identified in DDA (blue) and DIA (red) acquisition mode, respectively for decreasing loads of HEK293 tryptic peptides. The bars in the negative and positive directions represent the number of protein groups identified in common (intersection) or in total (union) for the technical triplicate injections at the indicated peptide loads, respectively. **c**. Distribution of the coefficients of variance (CV) for the peptide precursor intensities for the technical triplicate injections for each sample load. **d**. Distribution of the fold change (log2 scale) of the average precursor intensities between a given sample load and that at 2,000 ng sample load.

Next, we assessed the quantitative reproducibility and accuracy of the dilution series data generated in DIA mode. An average peptide quantification coefficient of variation (CV) of less than 10% was obtained for triplicate injections across the whole dilution series, even at a level of 3.9 ng (Fig. 1c). The run-to-run correlation coefficient was above 0.96 throughout the dilution series (Supplementary Fig. 4). The peptide quantification values obtained for the consecutive dilution steps retained linearity throughout the entire dilution range (Fig. 1d), and deviated by less than 20% in accuracy compared to the values at the highest sample load. It is noteworthy that for lower sample amounts (<125 ng) we observed a reduction in signal intensity for late eluting hydrophobic peptides (Supplementary Fig. 5), which we attribute to adsorption effects (e.g. to the walls of the sample tubes). To maintain sufficient robustness for the measurement of all peptides irrespective of their hydrophobicity, we therefore decided to set 100-200 ng of peptides as the practical lower limit for the subsequent measurements.

In summary, these results indicate that an optimal management of fill time on the Orbitrap Lumos significantly extends the performance of DIA proteomics towards sample amounts in the low ng range with minimum attrition in terms of identified proteins and quantitative accuracy.

### Optimization of sample handling for low cell numbers prior to MS analysis

To adapt the sample workup procedure for the low number of sorted cells required to yield the 100-200 ng of peptide mass per DIA measurement we devised a single tube procedure that minimized sample losses (Supplementary Fig. 6a). We identified several steps that were critical for the generation of high quality peptide samples, including: (i) FACS in the absence of fetal bovine serum (FBS), (ii) the use of low protein-binding plastic-ware to minimize the loss of hydrophobic peptides, (iii) careful pelleting of the non-adherent sorted cells, (iv) freeze-drying of the pellets and subsequent lysis and digestion in the smallest volume that can be handled practically (Supplementary Fig. 6b-c and expanded details in method section).

This optimized sample processing protocol was used to prepare triplicate samples of 200,000, 100,000, 50,000, 25,000 and 12,500 and 6,250 CD34+ cells obtained by FACS. We determined that ca. 3.2 μg of total peptide mass could be recovered from 200,000 sorted cells after the entire process of which approximately 73% (8/11 μl) could be injected by the autosampler for DIA measurements. By extrapolation, we therefore estimated that we injected approximately 2,327, 1,164, 582, 291, 145, and 73 ng of peptides, respectively, from the lower number of sorted cells. In comparison, 26.8 μg of peptide mass was obtained from 200,000 HEK293 cells when processed in bulk. This is in agreement with the 4-5 times smaller cell volume expected for the CD34+ cells compared to HEK293 cells ^24,25^.

The average number of identified protein groups decreased from 6,955 for 200,000 to 4,833 for 12,500 CD34+ sorted cells. A median CV below 14% (Fig. 2b, Supplementary Table 2) and a good linearity of quantification was also maintained for these measurements (Fig. 2c). At 6,250 cells, the number of identified protein groups decreased to 2,248 and the CV of quantification increased to above 17%. However, the overall peptide quantification remained relatively well correlated throughout the entire CD34+ series (Supplementary Fig. 7), indicating that protein quantification remained quite accurate even for samples containing as few as 6,250 cells. These 2,709 protein groups were almost entirely subsumed in the set of proteins identified at higher sample loads (Supplementary Fig. 8), indicating that meaningful comparisons across sample cohorts are feasible even in cases in which one or several samples are only available in minute quantities.

**Figure 2.**
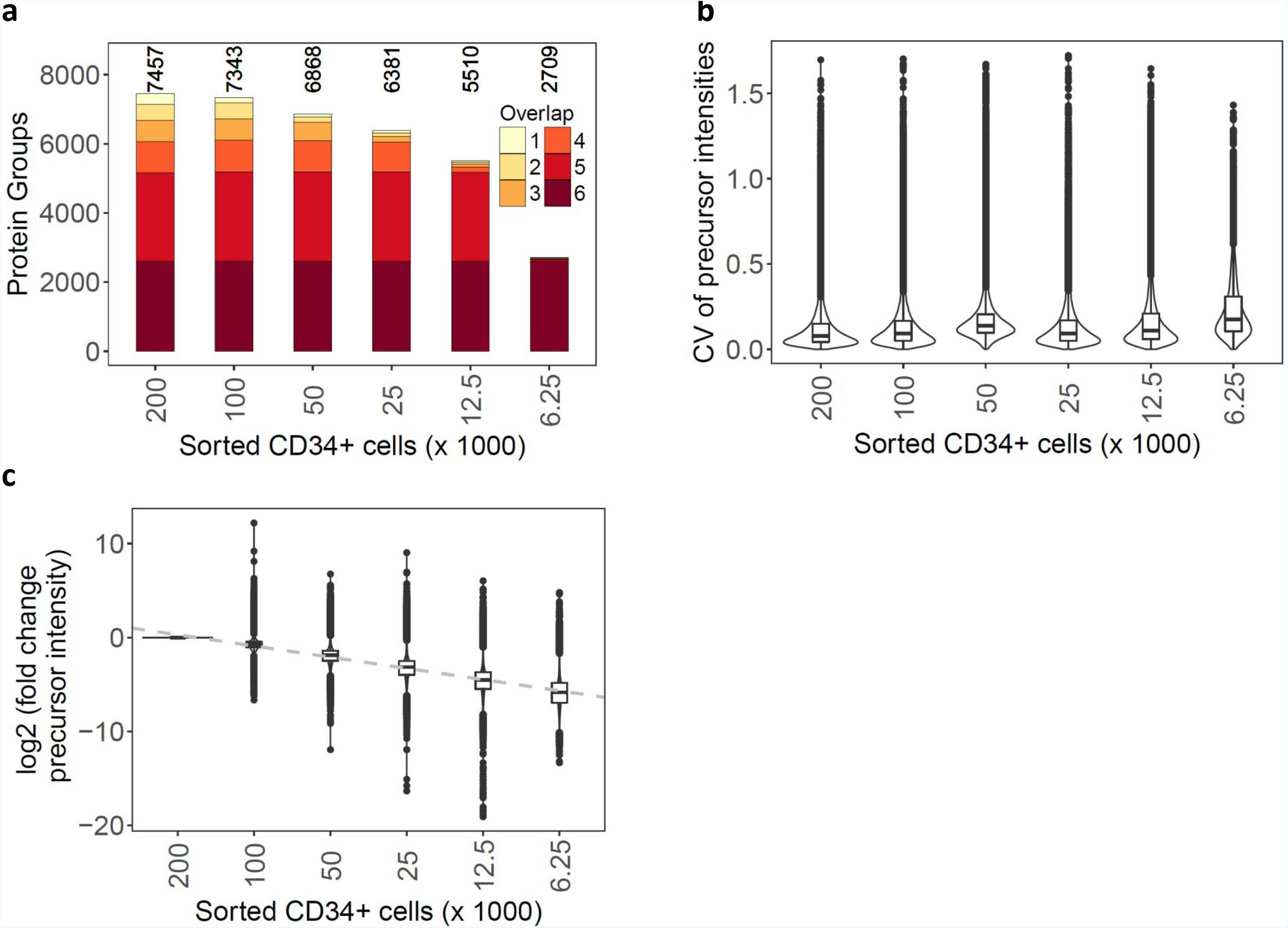
**a.** Number of protein groups cumulatively identified across the technical replicates for decreasing numbers of FACS-isolated CD34+ cells. The color scale represents the consistency of protein group identifications across the runs **b.** Distribution of the coefficients of variance (CV) for the peptide precursor intensities for the technical triplicate injections for FACS-isolated cells. **c**. Distribution of the fold change (in log2 scale) of the average precursor intensities between a given sample load and that at 200,000 FACS-isolated CD34+ cells.

Relating the results from sorted cells to those of the HEK293 peptide dilution series, we noted an attrition in the number of identified proteins and their quantification accuracy for cell numbers below 25,000, corresponding to <300 ng of peptide mass on column. Overall, the optimized DIA and sample preparation method provided reproducible identification and quantification results (in all three replicates) for more than 5,100 proteins from as little as 25,000 CD34+ FACS-isolated cells.

### Proteomic analysis of human hematopoietic cell subpopulations underscores ontogenetic distance between individual cell types

We applied the optimized method to profile the proteome of CD34+ cell subpopulations isolated from the peripheral blood of five HSPC donors (age 28-57 y, see Supplementary Table 3). Four highly enriched subpopulations (HSCs/MPPs, CMPs, MEPs and GMPs), respectively, were isolated by FACS, processed (for details, see Online Methods) and analyzed by DIA-MS (Fig. 3a and 3b). Guided by the cell dilution experiment described above, 25,000 cells were collected for each subpopulation. To support peptide-centric analysis of the DIA data, we generated a spectral library specific for the cell types of this study (see method section). We identified on average 5,851 protein groups for the different hematopoietic stem cell and progenitor cell populations (Supplementary Fig. 9, Supplementary Table 4). To increase the robustness of the following differential comparison, we applied additional stringent filtering criteria that resulted in a final list of 4,131 protein groups (from 39,264 peptide precursors) that were quantified consistently with at least two peptides across the samples.

**Figure 3.**
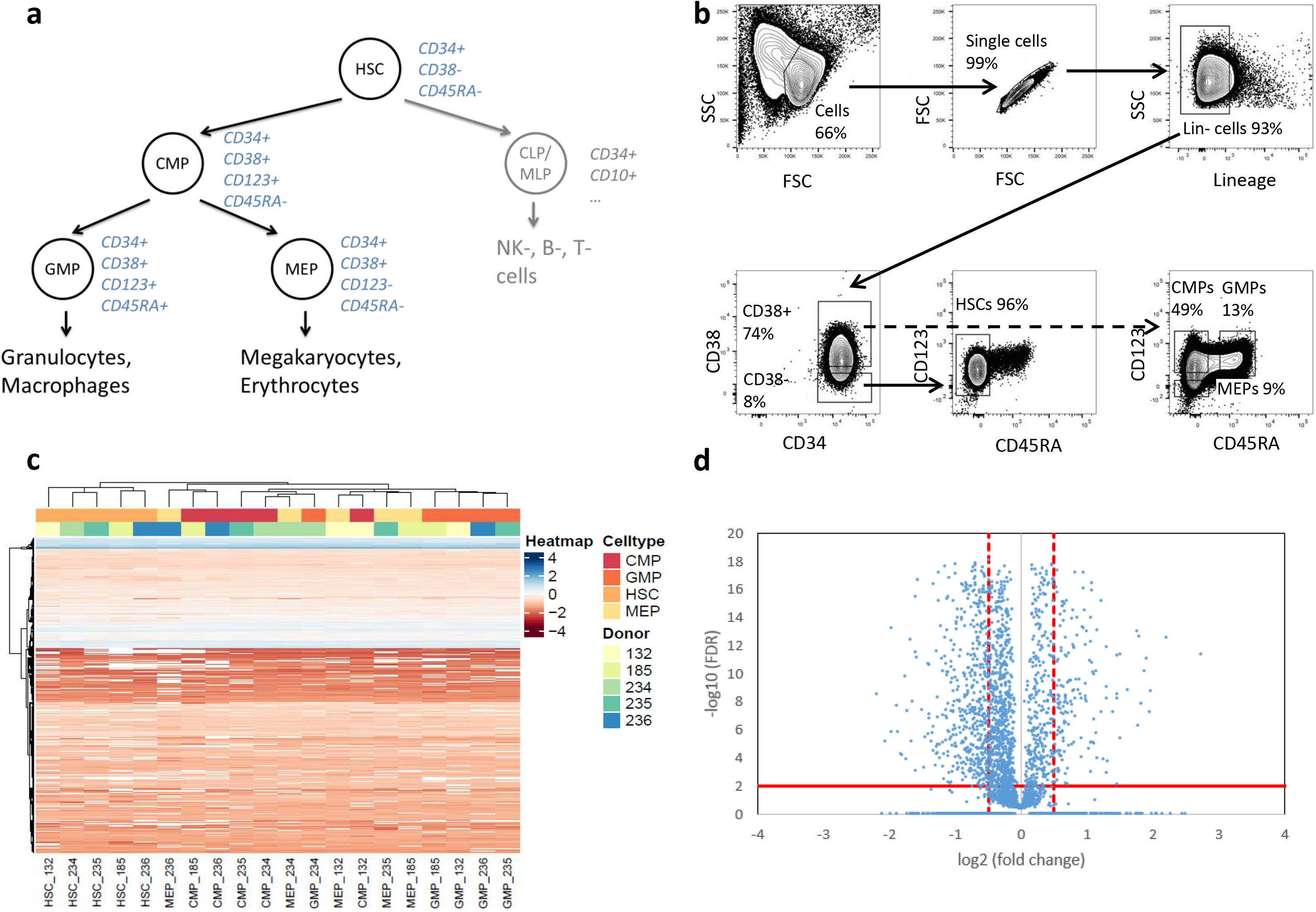
**a.** Human hematopoietic cell hierarchy with respective cell surface markers depicted in blue ^7-9^. **b.** Fluorescence-activated cell sorting (FACS) strategy, depicted on magnetic-activated cell sorting (MACS)-preselected CD34+ cells isolated from healthy HSPC donors. Shown are the analysis gates. Highly enriched HSCs/MPPs (referred to as HSCs) are CD34+CD38-CD45RA-, highly enriched CMPs are CD34+CD38+CD123+CD45RA-, highly enriched GMPs are CD34+CD38+CD123+CD45RA+ and highly enriched MEPs are CD34+CD38+CD123-CD45RA− **c.** Non-hierarchical clustering (Manhattan distance) heatmap of intensities for the peptides identified in HSCs, CMPs, GMPs, MEPs (shades of red) isolated from five different donors (shades of blue). The peptide intensities are centered and scaled and depicted in color shades from red to blue. The missing peptide intensity values are shown in white. **d.** Volcano plot of differential analysis of proteins. Comparison of HSCs to the average of the three other cell types. Abbreviations: HSPC – hematopoietic stem and progenitor cell; HSC - hematopoietic stem/multipotent progenitor cell; CMP - common myeloid progenitor; CLP/MLP - common/multipotent lymphoid progenitor; GMP - granulocyte-macrophage progenitor; MEP - megakaryocyte-erythrocyte progenitor; SSC - side scatter; FSC - forward scatter

Because the cell samples were derived from non-related donors of different age and the analyzed cell populations are relatively close in the cell differentiation tree, we first tested whether the quantitative protein measurements were sufficiently accurate and reproducible (see Supp. Fig. 10 and 11 for further details) to confidently detect cell subtype specific differences despite the expected inter-person variability. The summary heat map of these comparisons (Fig. 3c) indicates that the proteome profiles clustered according to cell subtype rather than donor. HSCs/MPPs clustered the furthest away from the other cell subpopulations, while CMPs were found to be more similar to GMPs and MEPs for some donors, in agreement with the ontogenetic distances expected between the different cell lineages (Fig. 3a).

Proteins identified as differentially expressed in the various cell subpopulations were distributed with homogenous abundance range with a significant proportion lying beyond the cut-offs of FDR <0.01 and log2(fold change)>0.5 (Fig. 3d and Supplementary Fig. 12). Using these cut-offs and comparing HSCs/MPPs to the average of the remaining three subpopulations, 1,008 proteins were determined to be differentially regulated in HSCs/MPPs. In accordance with the close ontogenetic distance of CMPs to GMPs and MEPs, the number of proteins with significantly changed expression was somewhat lower for GMPs (489) and MEPs (370) and even lower for CMPs (64).

Overall, these results show the consistent identification of quantitative protein patterns that are characteristic for ontogenetically close cell types, even within the genetic variability of a human population.

### Gene ontology enrichment analyses recapitulate the expected changes for proteomics and transcriptomics data

From the same sorting experiments, a further 10,000 cells were isolated for RNAseq. Similar to the proteomic results, the transcriptomic data also revealed clear clustering according to cell type rather than donor (Fig. 4a), with the HSCs/MPPs clustering furthest apart from the other cell subpopulations, while CMPs were more similar to GMPs and MEPs.

**Figure 4.**
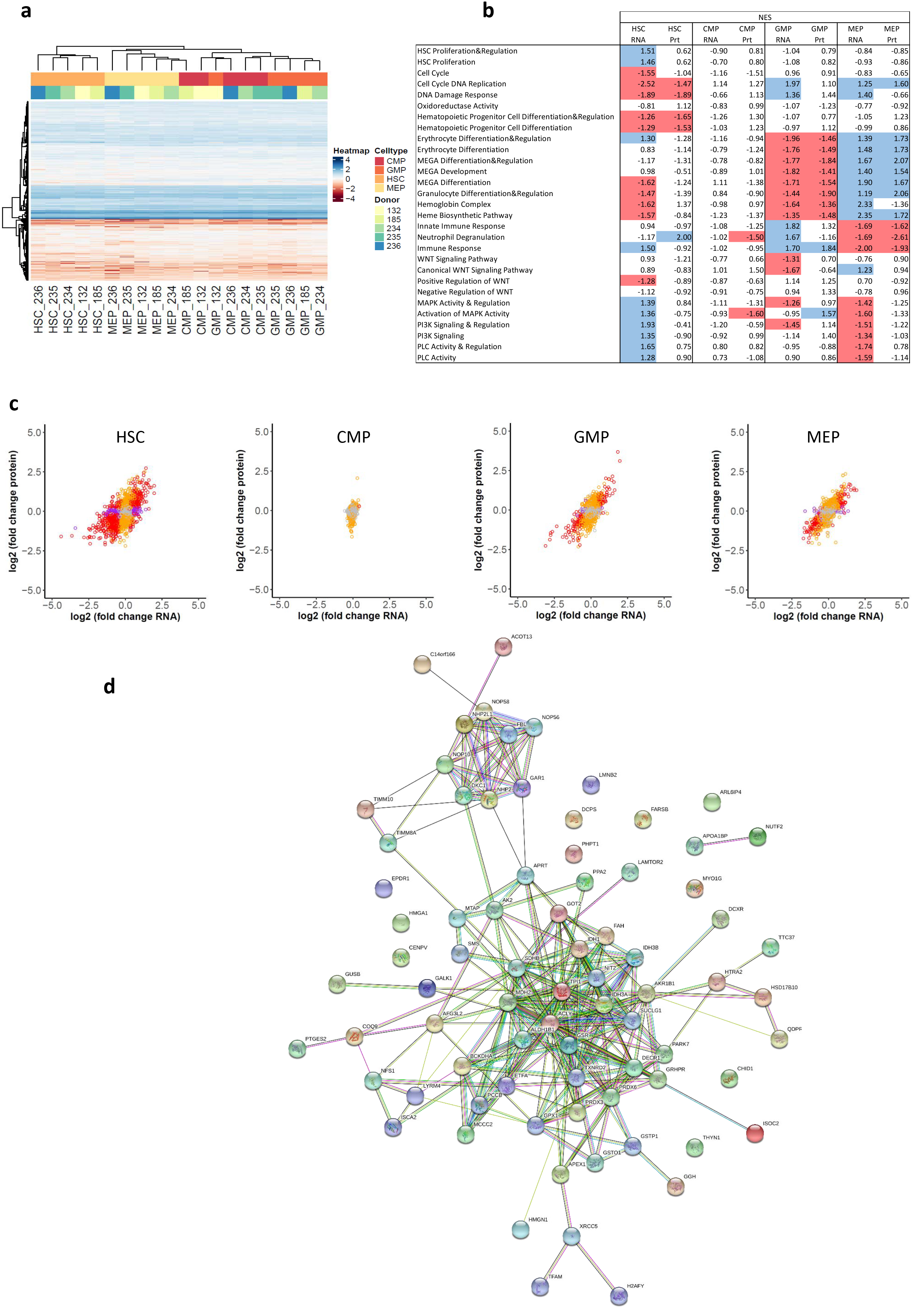
**a.** Non-hierarchical clustering (Manhattan distance) heatmap of the intensity for the transcripts identified in HSCs/MPPs (referred to as HSCs), CMPs, GMPs, MEPs (shades of red) isolated from five different HSPC donors (shades of blue). The transcript intensities are centered and scaled and depicted in color shades from red to blue. The missing transcript intensity values are removed because they could not be handled by the clustering algorithm. Clustering was observed according to cell type, not according to donor. **b.** Gene Ontology Enrichment Analysis showed good alignment of protein and mRNA data. Gene set enrichment analysis (GSEA) was performed for ranked mRNA and protein lists using GO processes from the Gene Ontology Consortium database as gene sets. Shown are normalized enrichment scores (NES) for the individual gene sets. Significantly upregulated gene sets are marked in blue color, significantly downregulated gene sets are marked in red color. Significance was defined as FDR < 0.25, specific cell subpopulations were compared to the average of the remaining three cell types, and log2(fold change) was used as ranking criterium. Abbreviations: MEGA – megakaryocyte; MAPK - mitogen-activated protein kinase; PI3K - phosphoinositide-3-kinase; PLC - phospholipase C. **c.** Correlation between proteomics and transcriptomics data for HSCs/MPPs (referred to as HSCs), CMPs, GMPs and MEPs. Dots are depicted in red when the FDR was below 0.01 both for protein and RNA data, orange when FDR < 0.01 for protein data, purple when FDR < 0.01 for RNA data < 0.01, and grey when FDR ≥ 0.01 for both protein and RNA data **d.** Network analysis of significantly upregulated proteins with concomitant significantly downregulated mRNA in HSCs/MPPs. Two clusters were especially prominent including the snoRNPs and telomerase complex proteins GAR1, DKC1, NOP10, NHP2 and the quiescence-inducing NAD(P)H-producing proteins IDH1, IDH3A, IDH3B. HSCs/MPPs were compared to the average of the other three subpopulations; cut-offs were set at FDR < 0.01 for protein and RNA data.

To further assess our proteomic and transcriptomic results, we performed gene set enrichment analyses for specific gene ontology (GO) processes involved in hematopoietic stem cell differentiation for the enriched cell subtypes according to previous studies and functional annotations ^1,2,10,11,16,26-28^. The transcriptomic and proteomic data recapitulated well most of the expected changes in GO processes (Fig. 4b and Supplementary Fig. 13, Supplementary Fig. 14). Cell cycle/DNA replication/DNA damage response was found downregulated for the overall more quiescent HSCs/MPPs both on mRNA and protein level. Erythrocyte differentiation/ megakaryocyte development and differentiation/heme biosynthesis were observed to be upregulated in MEPs on mRNA and protein level. (Innate) immune responses were found upregulated in GMPs on mRNA and protein level. The canonical WNT pathway was observed to be upregulated in MEPs on the mRNA level. Mitogen-activated protein kinase, phosphoinositide-3-kinase and phospholipase C pathways were all shown to be upregulated in HSCs at the mRNA level ^1,2,10,11,27,28^.

Of note, the transcriptomic and proteomic results showed highest agreement of the GO enrichments for the GMPs and MEPs, both in terms of directionality (up or down) and significance of the pathway enrichments. For the CMPs, only few significant pathway enrichments were observed, probably due to the composite nature of the CMPs between GMPs and MEPs. Interestingly, the HSCs/MPPs showed fewer significantly up- or downregulated gene ontology processes for the proteomics data compared to the transcriptomics results (e.g. HSC proliferation, MAPK activity and regulation, PI3K signaling, PLC activity, see Fig. 4b), highlighting the importance of identifying a large number of entities (close to 20,000 mRNAs versus a few thousand proteins) as well as the possibility of alternative methods of regulation for proteins (e.g. localization, post-translational modification etc.). For specific pathways (e.g. oxidoreductase activity, erythrocyte differentiation and regulation, neutrophil degranulation, see Fig. 4b), discordant directionality of enrichment between the proteomics and transcriptomics results was observed, despite not reaching significance, emphasizing the complementary value of the protein quantitative information for quiescent cells in which the transcription machinery is known to be strongly reduced.

Overall, the GO pathway analysis was in agreement with the expected properties of the respective cell types, thereby validating our mRNA and protein results against former studies.

### Discrepant mRNA and protein regulation in highly enriched HSCs/MPPs

To investigate the potential discrepancy between the RNA and protein patterns suggested by the GO enrichment analysis, we decided to examine the proteins that were found differentially regulated compared to their mRNAs in more detail (Fig. 4c).

The protein vs mRNA fold-change plots (Fig. 4c) showed good correlation for MEPs (R^2^ of 0.50) and GMPs (R^2^ of 0.41), but somewhat less for the HSCs/MPPs (R^2^ of 0.32) and CMPs (R^2^ of 0.06), in line with the results from GO enrichment analysis (Fig. 4b).

The STRING analysis of proteins differentially regulated at the transcript and protein level revealed two major protein-protein association networks, one including the small nucleolar ribonucleoproteins (snoRNPs) and telomerase maintenance proteins GAR1, DKC1, NOP10, NHP2, and the other the quiescence-inducing NAD(P)H-producing isocitrate dehydrogenase proteins IDH1, IDH3A, IDH3B (Fig. 4d, Online Methods). Both association networks were upregulated on protein and downregulated on mRNA level in HSCs/MPPs (see also Supplementary Fig. 15).

## DISCUSSION

One of the frontiers of modern biology lies in deciphering how individual cells operate in context. Typically, due to a lack of sufficiently sensitive analytical techniques, averaged measurements of pools of cells that are heterogeneous in cell type, localization or cell cycle state have been carried out. Nowadays, modern technologies promise to dissect the biochemical processes in play at the single cell level with unprecedented resolution. Extensive transcript data can indeed be readily obtained at the single cell level ^29^. This cannot be achieved yet at the proteome level, even though quantitative protein patterns should provide invaluable complementary information about the biochemical state of small cell populations, compared to transcript profiles ^30^.

Using characteristic markers for cellular differentiation, FACS is not only a tool for the analysis of individual cells, but also provides homogeneous, live, sorted cell subpopulations, thereby offering an ideal link between complex cell samples, their cell type composition and their respective biochemical/pathway and phenotypic analysis. In most cases, sorted cells from biologically relevant samples (e.g. patients) are only available in minute amounts and without replicates. To achieve the most comprehensive and accurate protein quantification possible for those precious samples, we adopted DIA acquisition on an Orbitrap instrument to maximize the sensitivity of the workflow.

We show that the external ion accumulation on such a platform is best exploited when using DIA at very low sample loads. Once the fill times approach the Orbitrap scan time, the degree of parallelization is maximized, and highly accurate and reproducible protein quantification even for sub-100 ng of total peptide mass loaded on column can be realized. Importantly, our results also demonstrate that even though the number of protein identifications decreased when lowering the amount of material injected, protein quantification retained a very high level of accuracy (Fig. 1d, Fig. 2c, Supplementary Fig. 4, Supplementary Fig. 7). In practice, this means that even if less than 25,000 cells are available from FACS, protein quantification results can still be confidently compared to those from higher cell counts, if the proteins are detectable in both samples.

We applied the developed DIA acquisition scheme to the analysis of 25,000 FACS-isolated human hematopoietic stem and progenitor cell (HSPC) subpopulations and could quantify more than 5,800 protein groups per cell subtype, of which we used a stringently filtered subset for further analysis. Proteome and corresponding transcriptome data showed the same clustering pattern. Interestingly, while the GO analysis showed high levels of agreement for the mRNA and protein data of GMPs and MEPs, there were discrepancies for the HSC/MPP subpopulation. Given the similar data quality (no. of protein groups identified, quantification, coefficient of variation) for the various analyzed cell types, the different behavior of HSCs/MPPs is likely to be biologically significant.

HSCs are mostly quiescent cells, i.e. with major fractions in G0 of cell cycle, which have been shown to contain very low levels of mRNA, while still maintaining a relatively constant protein mass overall. Proteins may therefore provide a better read-out of the cellular state of quiescent cells than the mRNAs. We therefore decided to focus our analysis on the proteins that showed discrepant regulations between their proteomic and transcriptomic data ^31^ and identified two strongly interconnected nodes which are upregulated on the protein level whilst downregulated on the mRNA level in HSCs/MPPs. The first protein-protein association network includes several snoRNPs and telomerase maintenance proteins deemed essential for long lived stem cells and demonstrated to be differentially regulated on protein and mRNA level ^32-34^. Telomerase activity in hematopoietic cells is associated with self-renewal potential and has been shown to decrease upon myeloid differentiation ^33^. Mutations in these telomerase maintenance proteins result in dyskeratosis congenita, a syndrome characterized by bone marrow failure and an increased risk for acute myeloid leukemia and myelodysplastic syndromes ^35^. Members of the second cluster include the isocitrate dehydrogenase (IDH) proteins IDH1, IDH3A and IDH3B that have previously been shown to maintain quiescence in hair follicle stem cells ^36^. IDHs are also thought to play a key role in hematopoietic stem cell homeostasis and were reported to be mutated in approximately 20% of acute myeloid leukemias ^37,38^. In line with those observations, low IDH mRNA levels had also been described in cultured hematopoietic stem cells ^39^.

These two examples illustrate the relevance of generating high quality proteomic data for well-defined cell subpopulations with the goal to identify biological processes that cannot be detected by genomic or transcriptomic analysis. Though this seems particularly evident for quiescent cells, we expect that proteomic data will bring an invaluable layer of biological information complementary to that of the transcriptomic data for many other cell subtypes. The presented methodology can thus be expected to increase our understanding of the dynamics of cell type-specific networks and to complete our knowledge on differentiation processes at play in healthy and pathological cells ^11,27^.

It will be important to further refine the different cell subpopulations. CD34+CD38-CD45RA− HSCs/MPPs, CD34+CD38+CD123+CD45RA− CMPs, CD34+CD38+CD123+CD45RA+ GMPs and CD34+CD38+CD123-CD45RA− MEPs could indeed be further divided into biologically even more refined subpopulations ^2,8^. Also, proteomic data from cell types isolated directly from bone marrow or from cord blood rather than from mobilized HSPCs obtained from donors after artificial stimulation will need to be obtained. For both cases, further fine-tuning of the sample handling steps may be necessary to cope with possibly lower sample amounts. This concerns, among other parameters, the observed adsorptive losses to surfaces. Possible avenues to explore would be the use of novel microfluidic devices ^40^ and the use of LC columns with further reduced inner diameters ^41^, strategies that have recently been used in combination with Orbitrap Lumos instruments operated in DDA mode. Thereby, the ultimate goal is the application of the proteomic profiling technology to clinically relevant rare cell types such as (pre)leukemic stem cells in hematopoietic malignancies as well as cancer stem cells from solid tumors.

In summary, we describe an ultra-sensitive mass spectrometric method that allows to generate highly accurate and reproducible protein quantification data from minute amounts of highly homogeneous cell subpopulations enriched by fluorescence-activated cell soring (FACS). This technology bridges the gap between the proteome profiling of samples containing multiple cell types and single cell analysis by allowing to dissect the biochemical processes in play in specific cell subpopulations of interest with unprecedented sensitivity, depth of coverage and reproducibility.

## METHODS

Methods and any associated references are available in the online version of the paper.

## ACKNOWLEDGMENTS

We are grateful to Audrey van Drogen (ETH Zurich) for providing an aliquot of HEK293 cells, Lukas Reiter and Oliver Bernhardt (Biognosys AG, Schlieren) for the Fill Time Reporter software to extract the fill times from the raw data files and Alexander Leitner (ETH Zurich) for support with MS method development and manuscript preparation. We thank Patrizia Belleda and Asuka Fry (University Hospital Zurich) for the initial processing of patient samples and Claudia Dumrese (University of Zurich) for assistance with FACS. We acknowledge Witold Wolski (Functional Genomics Center Zurich) for assistance with the statistical analysis of the omics data and data interpretation.

This study was supported by two grants to Fabienne Meier-Abt (Filling the Gap Grant of the University of Zurich; Krebsliga Zurich Grant) and the Clinical Research Priority Program Human Hemato-Lymphatic Diseases of the University of Zurich (Markus G. Manz). Alexandre P.A. Theocharides was supported by the Cloëtta foundation. Sabine Amon was supported by the ERC grant “HLA-DR15 in MS” (FP7-IDEAS-340733); Ludovic Gillet was supported by the grants “Proteomics 4D” (ERC-2014-AdG 670821) and “PrECISE” (H2020-EU.3.1 668858). Access to the instrumentation was supported by the following grants to Ruedi Aebersold: “Proteomics 4D” (ERC-2014-AdG 670821), “ULTRA-DD” (FP7-JTI 115766) and ETH Scientific Equipment.

## AUTHOR CONTRIBUTIONS

R.A., M.G.M. and A.P.A.T. conceived of the study and supported data analysis. S.A. and L.G. processed the HEK293 and HSPC samples. S.A. carried out the method optimization, the mass spectrometric measurements and the analysis of mass spectrometric data and differential analysis to the RNA data. F.M.A. prepared and FACS-isolated the HSPC samples, processed the samples for RNA sequencing, carried out the qPCR validation experiments and performed the comparative gene ontology/STRING analyses. S.D. performed mRNA quantification, normalization and differential expression analysis. S.A., F.M.A., L.G. and R.A. wrote the manuscript.

